# Sex biased expression and co-expression networks in development, using the hymenopteran *Nasonia vitripennis*

**DOI:** 10.1101/540336

**Authors:** Alfredo Rago, John (Jack) H Werren, John K Colbourne

## Abstract

Sexual dimorphism requires gene expression regulation in developing organisms. Differential expression, alternative splicing and transcript-transcript interactions all contribute to developmental differences between the sexes. However, few studies have described how these processes change across developmental stages, or how they interact to form co-expression networks. We compare the dynamics of all three regulatory processes in the sexual development of the model parasitoid wasp *Nasonia vitripennis*, a system that permits genome wide analysis of sex bias from early embryos to adults. We find relatively little sex-bias in embryos and larvae at the whole-gene level, but several sub-networks show sex-biased transcript-transcript interactions in early developmental stages. These provide new candidates for hymenopteran sex determination, including histone modification genes. In contrast, sex-bias in pupae and adults is driven by whole-gene differential expression. We observe sex-biased splicing consistently across development, but mostly in genes that are already biased at the whole-gene level. Finally, we discover that sex-biased networks are enriched by genes specific to the *Nasonia* clade, and that those genes possess the topological properties of key regulators. These findings suggest that regulators in sex-biased networks evolve more rapidly than regulators of other developmental networks.

## INTRODUCTION

Sexual differentiation during development involves many genes that are shared between sexes (i.e. genes on the autosomes) and therefore must be caused by epigenetic mechanisms of expression regulation. Indeed, gene expression regulation and alternate splicing are fundamental for sexual differentiation. For example, sexual development can be induced, even if the organism lacks the chromosomes normally associated with that sex (Capel 2017; Parma, Veyrunes, and Pailhoux 2016). Induction of a specific sexual phenotype can be achieved in the laboratory, either by changing the expression of sex-determining genes (Koopman et al. 1991; Colvin et al. 2001; Ventura et al. 2012; Verhulst, Beukeboom, and van de Zande 2010) or by administering hormones in the right developmental phase (Chew and Renfree 2016; Flament 2016; Olmstead and LeBlanc 2007).

In nature, numerous organisms lack sex-chromosomes altogether, demonstrating that sex-limited genes (i.e. genes located on sex-specific chromosomes) are not necessary for the evolution of phenotypic differences between sexes (Ge et al. 2018; Capel 2017; Holleley et al. 2016; Ashman et al. 2014; Verhulst, van de Zande, and Beukeboom 2010). Rather, phenotypic differences between sexes seem to evolve whenever the selective pressures between the two are different (sexually antagonistic selection or conflict, see Van Doorn 2009; Bonduriansky and Chenoweth 2009; Chippindale, Gibson, and Rice 2001; Kulmuni and Pamilo 2014). These results the same developmental components are often involved in the formation of different sexual traits, limiting the potential to optimize either sex independently (pleiotropic constraint). Characterizing the development of sexually dimorphic phenotypes is thus necessary to understand the regulatory variation that enables phenotypic diversification when male and female evolution conflict with each other.

Sex-biased gene expression is by far the most studied among the mechanisms that generate differences between sexes. Perhaps unsurprisingly, sex-biased expression of genes can account for up to 90% of all expressed genes in some species (Ingleby, Flis, and Morrow 2015). Most studies so far focused on measuring differences in the magnitude of gene expression only between adult males and females, and primarily in organisms with sex chromosome based sex determination (Chang et al. 2011; Innocenti and Morrow 2010 but see Wang, Werren and Clark 2015). However, the focus on adult gene expression does not reveal sex differential processes during development, and several genes are sex-biased only during those early stages (Perry, Harrison, and Mank 2014; Mank et al. 2010; Zhao et al. 2011). Changes in sex-bias expression of the same gene across development are most likely to create conflict, since the same locus may be under selection for changes in female-specific and male-specific functions in different developmental stages – a scenario that we term ‘developmental sexual conflict’. Several studies have also shown that alternative splicing can show sex-bias (Telonis-Scott et al. 2008; Hartmann et al. 2011; Brown et al. 2014), and that sex-biased splicing affects sex determination (Verhulst et al. 2013; Verhulst, Beukeboom, and van de Zande 2010) and sexual development (Chang et al. 2011).

The co-expression of multiple genes can result in effects that are qualitatively and quantitatively different from the sum of the effects of each gene (Ament et al. 2012; Spitz and Furlong 2012; Boyle et al. 2014), and provides a largely unexplored mechanism for the regulation of sex-specific differences; the same group of genes can have identical expression levels in males and females, yet still cause a sex-specific effect if they are only co-expressed in one sex (Arnold, van Nas and Lusis 2009; Van Nas et al. 2009). These interactions are therefore impossible to detect by independently testing transcripts, but can be identified *via* differential correlation analyses on transcriptional modules, or differential cluster correlations (Tesson, Breitling and Jansen 2010; Yang et al. 2013).

Differential cluster correlation (DC, right panel on Figure 1) is different from (and complimentary to) differential cluster expression (DE, left panel on Figure 1). Differential cluster expression measures the average fold expression change of a group of genes, and is thus analogous to gene-level differential expression (Langfelder and Horvath 2008). By contrast, differential cluster correlation measures the proportion of sex-specific interactions within the cluster (Zampieri, Soranzo and Altafini 2008; de la Fuente 2010). Since interactions require multiple genes, differential cluster correlation has no gene-level analog and is an emergent property of the transcriptional cluster. At the time of writing, few studies have attempted to systematically measure differential cluster correlations between sexes (Van Nas et al. 2009; Mozhui et al. 2012). As such, their importance is unknown.

**Figure 1:**
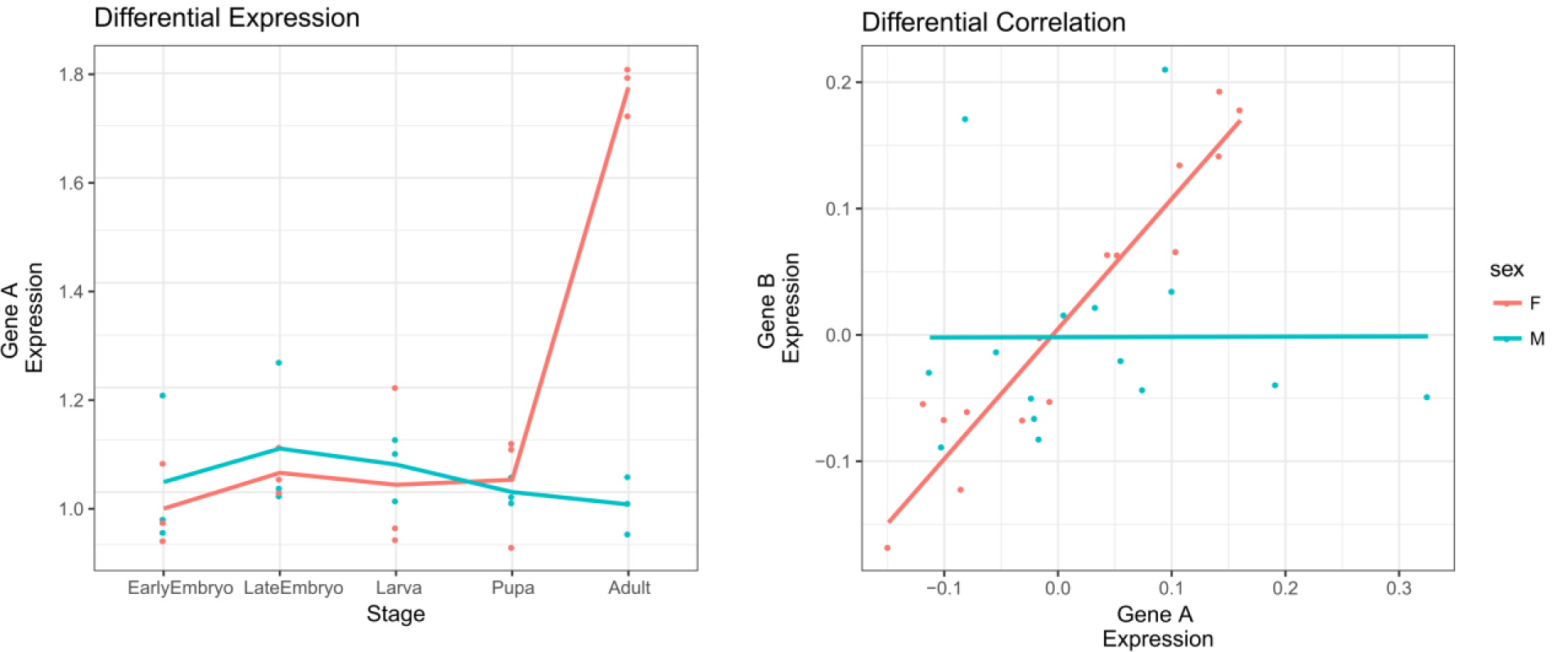
Examples of Differentially Expressed (left) and Differentially Correlated (right) clusters. Each dot represents a biological replicate of a gene’s expression value. In the left panel, the expression levels of the same gene are compared between sexes at each stage. DE is present if the gene expression values between the sexes differ. In the right panel, the expression levels of two genes are compared across samples for all stages. DC is present if the correlation between the two genes differs between the sexes.

We describe the developmental dynamics of these three forms of gene regulation (expression, splicing and co-expression) in the development of the jewel wasp *Nasonia vitripennis*. *Nasonia* has several advantages for investigating sex differential gene expression across development. First, like other hymenopterans (ants, bees, and wasps) *Nasonia* has haplodiploid sex determination (Whiting 1968): males and females have the same genomes (there are no sex chromosomes) and sex differentiation is due to gene expression changes in the identical set of genes shared by the two sexes. Second, *Nasonia* can be readily inbred to produce nearly completely isogenic lines, further reducing genetic differences between individuals (Heimpel and de Boer 2008; Godfray 2010). Finally, haplodiploidy also allows the collection of exclusive male and nearly exclusive female samples during the early stages of development, since (under our experimental conditions) virgin females produce only sons and mated females produce ~85-100% daughters. We can thus characterize the differences in gene expression between sexes before the onset of phenotypic differences (Pultz and Leaf 2003).

*Nasonia* belongs to the “megadiverse” hymenopteran superfamily Chalcidoidea (henceforth referred to as “chalcids”), which is estimated to contain approximately 500,000 species (J. Heraty 2009; J. M. Heraty et al. 2013). Most chalcids are small parasitoid wasps that prey on other arthropods, and play vital roles in regulating natural and agricultural ecosystems. *Nasonia* is used extensively in genetic, development, and evolutionary research. It’s genome was the first hymenopteran to be sequenced (Werren et al 2010) after that of the Honeybee (The Honeybee Genome Sequencing Consortium 2006). Several other hymenoptera genomes have been sequenced since (Branstetter et al. 2018), enabling us to study their genomic features in a broader phylogenetic context.

We utilize the *Nasonia* model system to investigate genome-wide sex differences in gene expression, differential splicing, and differential co-expression of genes across development from early embryonic stage to adulthood. We provide a first genome-wide characterization of *Nasonia*’s sex-specific transcript-transcript interactions via co-expression analysis across development. Finally, we examine the phylogenetic age of genes involved in these networks to investigate how they have evolved.

## RESULTS

### Few Loci Show Developmental Sexual Conflict

Across the five developmental stages (early embryo, late embryo, larva, pupa, adult), 6,041 out of 14,149 genes (43%) display sex-biased expression or splicing when tested individually (**Table 1**). Male-biased genes are prevalent at the pupal stage, whereas female-biased genes are prevalent at the adult stage. This pattern reflects the fact that spermatogenesis occurs primarily during the pupal stage and oogenesis primarily in adults (Clark et al. 2010). Larvae show the least amount of sex-biased transcription. Only one transcript (Nasvi2EG005321 or *Feminizer*) is sex-biased across all of development, followed by *Doublesex* (Nasvi2EG010980), which is female-biased in all stages from late embryo onward (>18 hours old). The low number of transcripts that are consistently differentially expressed across multiple stages is most likely due to the low number of sex-biased events in the pre-pupal stages. Only 751 out of 36,505 transcripts (2% of all transcripts) show sex-bias in the embryonic or larval stages: 203 occur in early embryos, 422 in late embryos and 154 in larvae. Transcripts that are differentially expressed in the embryonic stages are likely to be involved in sex-determination and will be of special interest for future studies.

**Table 1:** Number of sex-biased genes and transcripts at each developmental stage. Genes are counted as sex-biased if at least one of their transcription or splicing nodes is sex-biased.

| Stage: | Sex Biased Genes |  | Sex-biased Transcripts |  |
| --- | --- | --- | --- | --- |
|  | Male | Female | Male | Female |
| Embryo, 10 hr | 26 | 145 | 29 | 174 |
| Embryo, 18 hr | 187 | 185 | 202 | 220 |
| Larva, 51 hr | 17 | 121 | 17 | 137 |
| Pupa, yellow | 1,392 | 434 | 2,779 | 581 |
| Adult | 3,194 | 3,093 | 5,167 | 5,953 |

Transcripts that shift between male to female-bias in different developmental stages are considerably less frequent than expected by chance (Fisher’s exact test, p-value ~0); only 508 transcripts, generated by 373 genes show this pattern (6% of all sex-biased genes in our final dataset). The majority (66%) of these transcripts display shifts from male-bias in pupae to female-bias in adults. Other patterns that include both male and female-bias across development consist of changes from female-bias in various pre-adult stages to male-bias in adults (female-bias in pupa: 57 transcripts, female-bias in larva: 23 transcripts, female-bias in late embryo: 27 transcripts, and female-bias in early embryo: 8 transcripts). Interestingly, transcripts with pre-pupal sex-bias are significantly more likely to show shifts between male and female sex-bias than transcripts with post-pupal bias only (Fisher’s exact test, p-value ~0). All transcripts described in this study and their annotations can be found in Supplementary File S1.

### Low Relative Prevalence of Sex-Biased Splicing

Genes with sex-biased transcription are ~50% more frequent than genes with sex-biased splicing (6,041 versus 3,944). Over 67% of genes with sex-biased splicing also show sex-biased transcription, whereas less than 44% of genes with sex-biased splicing are also subject to sex-biased transcription (Figure 2). Only 1,294 genes show sex-biased splicing alone, compared with 3,391 genes with only sex-biased transcription. Taken together, these observations indicate that transcriptional bias is the main determinant of transcriptome-wide differences between the sexes. Our estimates on the fraction of sex-biased adult *Nasonia* genes are consistent with those previously reported (Wang, Werren, and Clark 2015).

**Figure 2:**
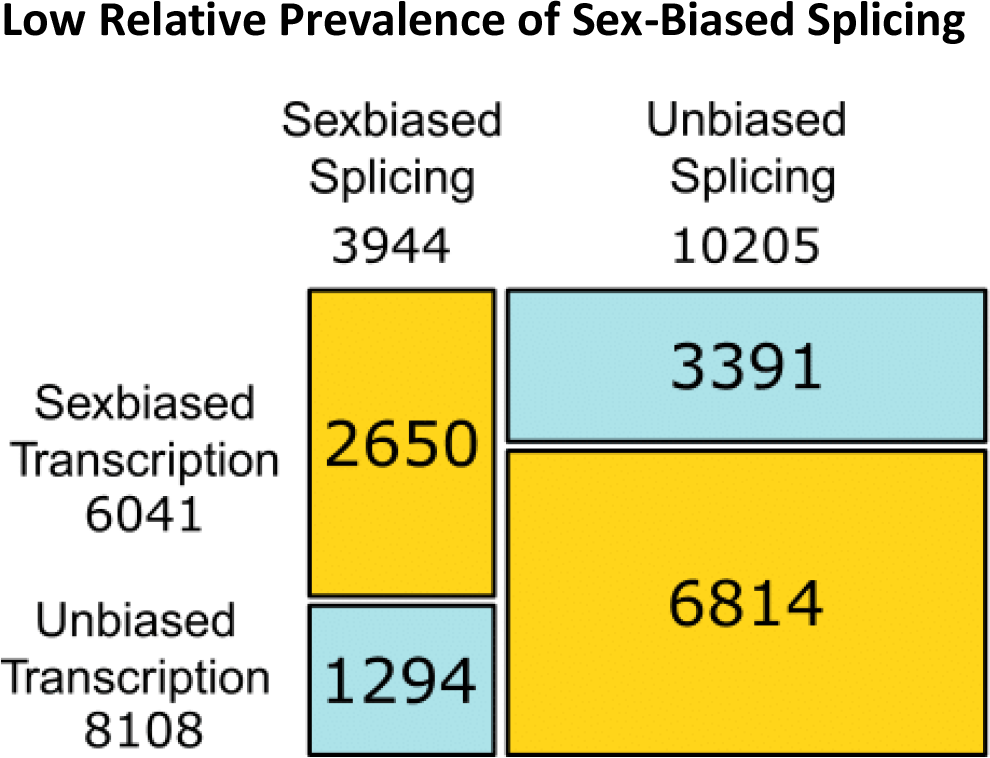
Proportion of genes with sex-biased splicing versus transcription. The area of each rectangle is proportional to the total number of genes with the corresponding combination of sex-biased splicing (x axis) and transcription (y axis). Numbers below each axis label report the total number of genes with sex-biased splicing or transcription and numbers in each square report the absolute number of genes in each combination. Yellow cells indicate over-represented combinations, and blue cells under-represented combinations.

### Few Genomic Regions Enriched by Sex-Biased Genes

We investigated whether regions of linked sex-biased loci are present in the *Nasonia* genome by searching chromosomal regions (Desjardins et al. 2013) for enrichment in male or female-biased genes. Current theories show that linkage could facilitate the evolution of adaptive sex-specific phenotypes by reducing the odds that alleles which work together are broken down by recombination (Thompson and Jiggins 2014).

Regions 1.065 and 5.072 show an enrichment for female-biased genes while region 4.1 is enriched in male-biased genes (Table 2). Female-biased group 1.065 contains *Nasonia*’s sex-lethal (Nasvi2EG000104) and histone deacetylase 3 (Nasvi2EG000106), a key component of histone-mediated gene regulation.

**Table 2:** Linkage groups enriched by sex-biased genes. Numbers indicate gene counts with their percentages compared to all genes in the linkage group. Recombination rates are expressed as centiMorgan per Mb. The last row reports median proportions and recombination rate across all linkage groups.

| Linkage Group | Enrichment | Male-biased Genes |  | Female-biased Genes |  | Total Genes | Recombination Rate |
| --- | --- | --- | --- | --- | --- | --- | --- |
| 4.1 | Male | 49 | 45% | 12 | 11% | 109 | 0.093 |
| 1.065 | Female | 3 | 8.3% | 18 | 50% | 36 | 0.38 |
| 5.072 | Female | 12 | 8% | 59 | 40% | 147 | 0.086 |
| Median Values | None |  | 17% |  | 17% |  | 0.25 |

Female biased linkage group 5.072 is strongly enriched by the controlled Gene Ontology (GO) annotation terms “apoptosis of nurse cells” (GO:0045476) and several other developmental terms related to photoreceptor and neuronal development (R3, R4 and R7 cell development and brain morphogenesis). Most genes within the male-biased linkage group 4.1 belong to cysteine-rich secretory proteins (PF00188.21). While these proteins are currently annotated as venom allergens, we hypothesize that the same secretory domains may, in this case, be involved in sperm production, as is suggested by expression patterns of their homologs in *Drosophila* (Kovalick and Griffin 2005).

Overall, the male enriched linkage group accounts for 1.2% of male-biased genes. The female-enriched linkage groups account for 2.0% of female-biased genes. While theory predicts selection for lower recombination rates in sex-biased genomic regions (Ellegren and Parsch 2007; Yeaman 2013), recombination rates in all three linkage groups fall within the interquartile range of recombination rates of all linkage groups.

### Clusters with Developmental Sexual Conflict Include Meiosis Genes

To provide a better overview of gene expression patterns, we grouped transcripts in co-expressed transcriptional clusters (henceforth **clusters**) and tested their average expression for significant sex-biased differential expression (**DE**). The results of analyses at the cluster level are quantitatively similar to the results obtained by examining single-nodes independently (Figure 3). Almost half of all clusters (81 out of 172) show DE at some point in development. Most DE occurs in pupal and adult stages (73 differentially expressed clusters), whereas only 8 clusters show DE in pre-pupal stages. The full assignments of transcripts to clusters is reported in Supplementary File S1, and the results of cluster-level analyses are reported in Supplementary File S2. Clusters are labeled as colour names, by convention (Langfelder and Horvath 2008).

**Figure 3:**
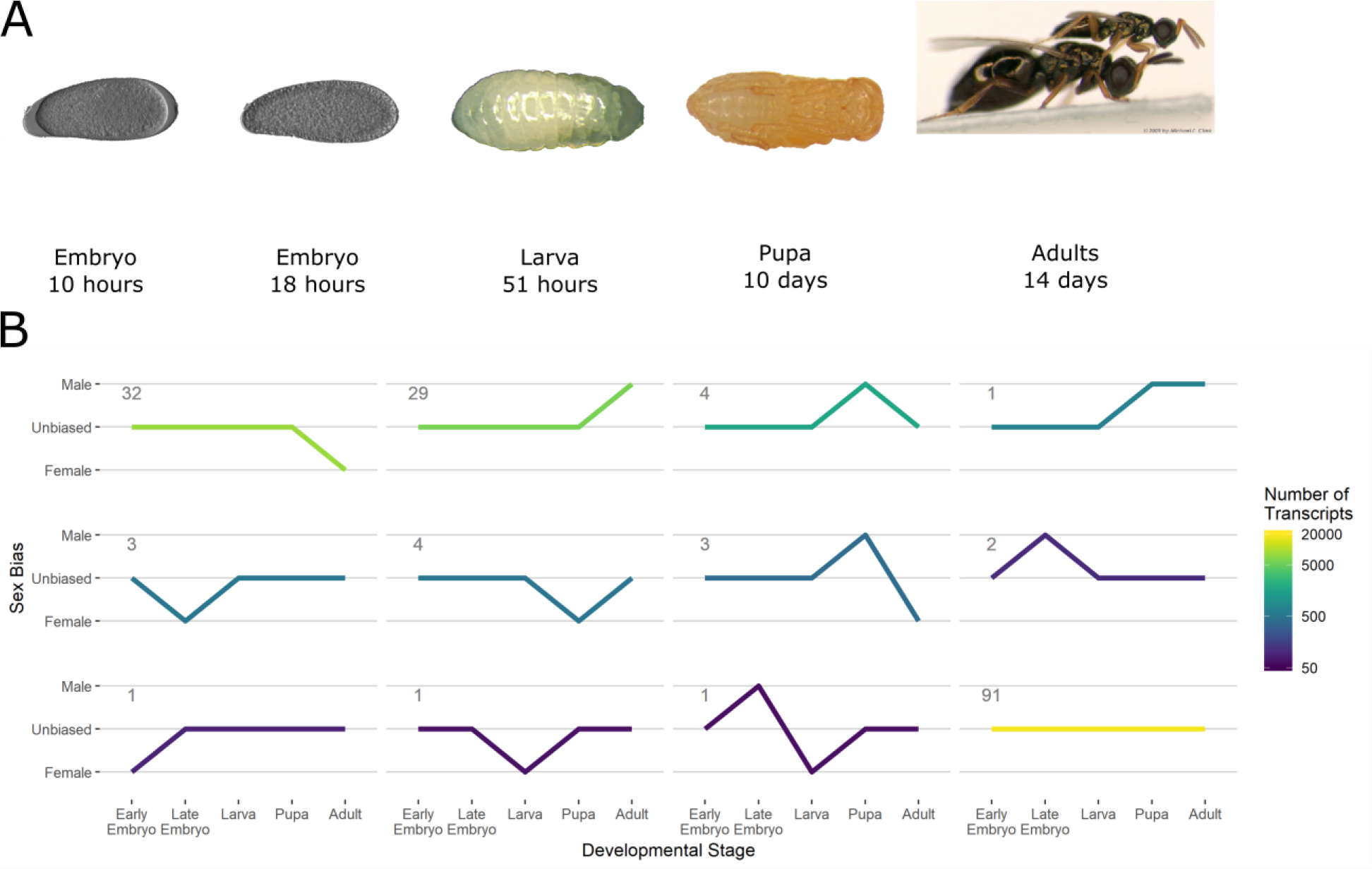
Differential expression (DE) of transcriptional clusters across developmental time from embryo (left) to adult (right). (A) Images of Nasonia at each stage sampled. (B) Patterns of sex-bias in each developmental stage. Each line represents a group of clusters which shares the same pattern of sex-bias according to GLMMs. The number of clusters that share the expression pattern is shown in the top left of each line. The total number of transcripts included in each clusters is represented by the colour (lighter for more transcripts, darker for less).

Four clusters alternate between male and female-biased expression in different developmental stages. Cluster ‘green3’ shifts from being male-bias in late embryos to female-bias in larvae. Its members consist of retro-transcriptases and unannotated multi-copy genes. It is therefore most likely due to transposon-related activity in the germline rather than developmentally related somatic processes.

The remaining three clusters (‘antiquewhite4’, ‘lightpink2’ and ‘yellow4’) shift from being male-biased in pupae to female-bias in adults. ‘Antiquewhite4’ and ‘yellow4’ contain multiple isoforms of the *Nasonia* SAK homolog (Nasvi2EG010310), a gene involved in the formation of sperm anoxeme in *Drosophila* (Bettencourt-Dias et al. 2005) and Cyclin B (Nasvi2EG014042), which is a conserved cell-cycle regulator that triggers mitotic division. Both clusters are enriched by meiosis and gametogenesis related GO annotations. Cluster ‘lightpink2’ contains several genes coding for amino acid binding proteins, including condensin (Nasvi2EG004100), which is involved in chromosome assembly and segregation (Hirano 2014).

Since *Nasonia* male gametogenesis occurs during the pupal stage, while female gametogenesis occurs primarily during adulthood (Clark et al. 2010), the shift in sex-bias observed in these clusters is likely caused by differences in the timing (heterochrony) of activation for gametogenesis-related genes. We further investigated the top-ranking hubs for each of those clusters, which mapped to three Constitutively Co-expressed Regulatory Events (CCRE 226, 345 and 3023 respectively). Each CCRE includes several transcripts with functionally identical expression profiles (see Materials and Methods for details, and Supplementary File S3 for full annotation of CCREs). When we compared the transcripts contained in each of those hubs with their Wasp Atlas entries (Davies and Tauber 2015), we discovered that other studies reported these to be moderately to extremely testes-biased in *Nasonia* (Akbari et al. 2013) thereby supporting the hypothesis that these clusters are primarily involved in male gametogenesis.

### Differential Correlation Reveals Early Sex-Biased Transcription

Our differential correlation (**DC**) analyses detected clusters that contain sex-biased transcript-transcript interactions (co-expression), regardless of whether the cluster is also differentially expressed (**DE**) between the sexes (see Materials and Methods for details). This analysis revealed 65 clusters of differentially correlated genes. Of those, 57 show differential correlation at the same time as differential expression (Figure 4); 7 out of the 8 clusters that show differential correlation in the absence of differential expression do so during the earliest stages of development. In other words, a majority of early transcriptomic differences between the sexes are revealed only when considering transcript-transcript interactions rather than individual gene expression levels. Sex-biased DC differs from DE in both timing and direction of change (Figure 4). No single cluster shows significant sex-biased DC in more than one stage; 4 clusters show DC in the earliest stages, and 5 out of the 65 DE clusters towards one sex in adults show an opposite bias in their DC. These initial observations suggest a degree of independence between DC and DE.

**Figure 4:**
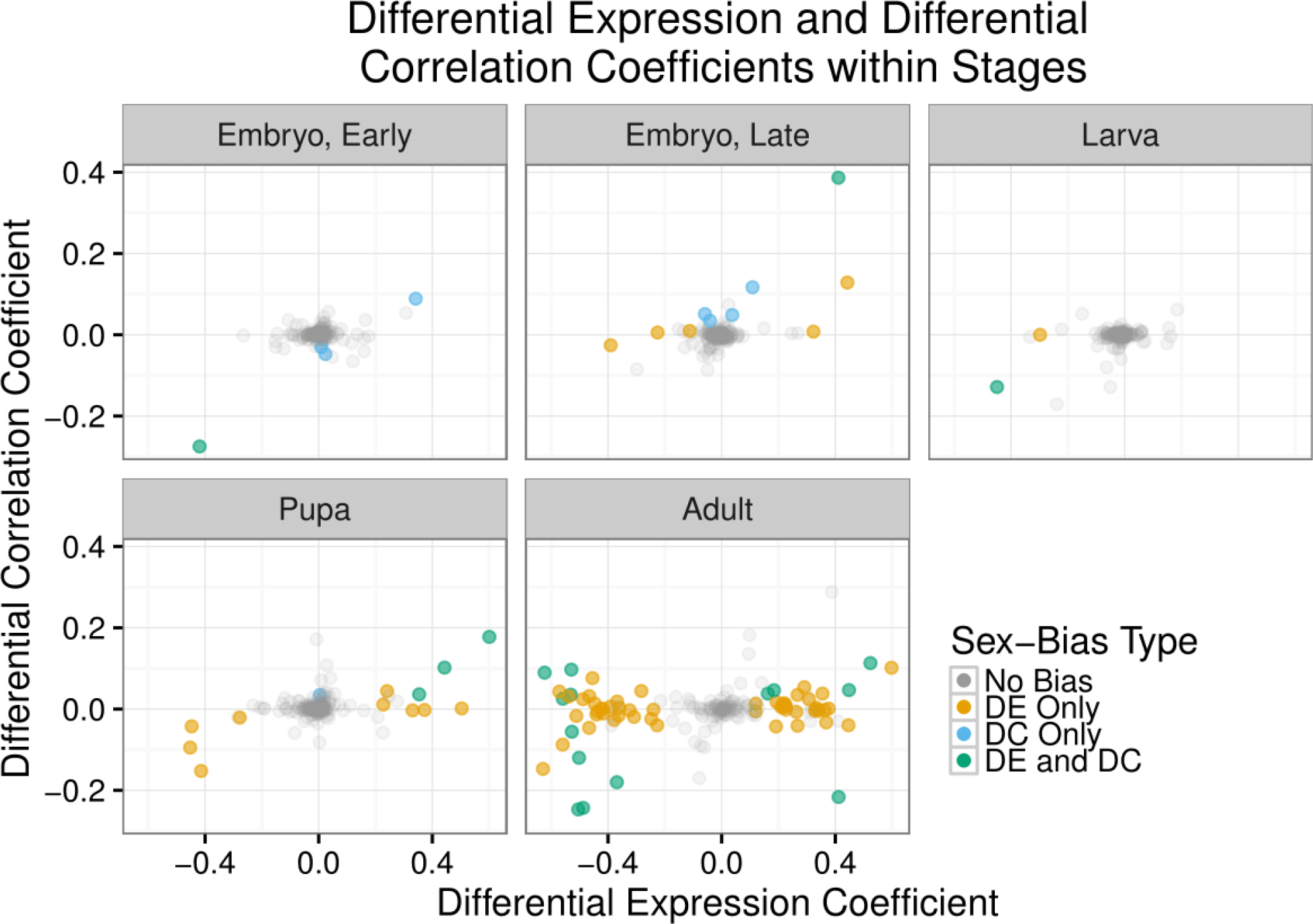
Cluster-level sex-biased expression and correlation across development. Each dot represents a single cluster in a single developmental stage. Position indicates the degree of sex-biased expression (x axis) and correlation (y axis). Positive values indicate male-bias, negative values indicate female-bias. Clusters are coloured in accordance with significant differential expression or differential correlation calculated from GLMs.

Only one of the DC clusters in early embryos is also DE (cluster ‘navajowhite3’), showing increased expression in female early embryos. This cluster is enriched by several GO annotations related to nucleosome assembly, comprising primarily of modified histone genes and possibly including the histone acetyltransferase complex H4/H2A HAT (genes Nasvi2EG008990, Nasvi2EG008772, Nasvi2EG024702, and Nasvi2EG008770). One of those genes is assigned to histone H1, one to histone H2A and two to histone H2B. These histone H2A and H2B nodes currently lack sufficient homology to be assigned to an orthologous group and are likely to be modified along a lineage-specific expansion of its gene family (Rago et al. 2016). Histone H1 is part of the most likely hub of this cluster (CCRE108), alongside an isoform of sex-lethal interactor (Nasvi2EG016490), and bällchen (Nasvi2EG003614) whose *Drosophila* ortholog is involved in the maintenance of neuronal and germline stem-cells via histone phosphorylation (Herzig et al. 2014; Yakulov et al. 2014). We therefore speculate that this cluster could be involved in sex differences mediated through histone modifications or chromatin condensation, and that genes in the cluster could be of special interest to future studies on the epigenetic mechanisms involved in sex-determination.

Two additional clusters (‘lavenderblush3’ and ‘palevioletred2’) do not show DE in early embryogenesis, but do show female-biased DC at that stage. Both are also differentially under-expressed in adult females relative to males. Why this pattern occurs is unclear but could be indicative of early sex determination processes. Neither cluster shows enrichment in informative GO annotations. CCRE 493 is the most hub-like node in cluster ‘lavenderblush3’, and is composed of the transcriptional nodes of gene Nasvi2EG018256 (a CDK inhibitor enriched in *Nasonia* testes relative to male carcass; Akbari et al. 2013), *Nasonia*’s Yellow-f protein (Nasvi2EG033442) and Inositol-trisphosphate 3-kinase A (Nasvi2EG003903), whose *Drosophila* homolog is necessary for correct wing formation (Dean et al. 2016). The primary hub of cluster ‘palevioletred2’ is CCRE 180, which groups two poorly characterized genes (Nasvi2EG007678 and Nasvi2EG001109), both of which are enriched in *Nasonia* testes relative to male carcass (Akbari et al. 2013). The same cluster also includes two fatty acyl-CoA reductases (Nasvi2EG017071 and Nasvi2EG025693) homologous to *Drosophila* and *Culex* male sterility proteins. Taken together, these observations suggest that these clusters could be involved in spermatogenesis, but cannot explain why they also show DC in early female embryos.

Notably, only one cluster (darkseagreen2) shows significant male-biased DC in early embryos. This cluster contains 75 genes and is strongly enriched by GO annotations related to stem-cell fate determination, neurogenesis and down-regulation of RNAs. Its hub node CCRE 1500 contains several genes which did not have orthologs with known functions in other organisms. In addition, the hub node includes a splicing event for Nasvi2EG022761: a homeobox-like transcription factor and Nasvi2EG006781, isoform of a testis-biased putative telomerase. We are uncertain about the potential role of these genes in very early sex differentiation, but they represent reasonable candidates for sex differences in early development. It should be noted in this regard that, in hymenopterans, males are derived from unfertilized eggs, and therefore likely require induction of specific pathways for development in early stage embryos.

While the direction of sex-bias is generally consistent between differential correlation and differential expression, we find that 5 of the 20 clusters with simultaneous DE and DC show bias in opposite directions. All of those exceptions are observed in adults. Four of these clusters (‘antiquewhite4’, ‘plum’, ‘plum3’ and ‘thistle3’) are more highly expressed in adult females yet more strongly correlated in adult males. The fifth (‘antiquewhite2’) is more highly expressed in males yet more strongly correlated in females. These discrepancies could be caused by differential tissue representation between the adult phenotypes, since females possess much larger gonads than males as a proportion to their body, and male spermatogenesis occurs primarily in pupae. The increased proportion of gonadal tissue could increase representation of non-sex-specific and male-biased gonadal transcripts in females, as well as their average expression compared to male gonadal transcripts. An increase in mean representation would affect differential expression analyses but not correlation-based ones, which rely on the relative change of transcript-transcript expression. This may be the case for cluster ‘antiquewhite4’, which (as mentioned earlier) is likely to be involved in gametogenesis. We also observe an enrichment for gametogenesis, neurogenesis, and histone modification associated annotations in the cluster ‘plum3’, while the cluster ‘thistle3’ is enriched by GO annotations related to germ cell development and splicing regulation. All genes contained in the hub nodes of those clusters show moderate testes-bias in adults of *Nasonia* (Akbari et al. 2013). The cluster ‘plum’ does not show significant enrichment in gametogenesis-related annotations, but rather is enriched by ribosomal biogenesis and RNA-processing related annotations. Both genes within its hub (CCRE106) are testes-biased (Akbari et al. 2013), suggesting involvement in either spermatogenesis or testicular functioning. By contrast, cluster ‘antiquewhite2’ is enriched mostly by GO annotations related to signal transduction and its hub contains several isoforms of Nasvi2EG010141, a calcitonin receptor enriched in female heads relative to bodies (Hoedjes et al. 2015). We surmise that this cluster may be involved in female-biased neuronal functioning and its apparent over-expression in males may be due to the relative smaller size of female brains compared to their gonads. Resolving some of these issues will require additional organ and tissue-specific analyses.

### Sex-Biased Clusters Have Different Regulatory Structure

To characterize network architecture of each cluster, we measured several of their topological and biological parameters (reported in Material and Methods). Since several of our measures showed non-trivial interactions (see Figure 9), we performed principal component analysis (PCA) to separate the combinations of features (Principal Components, or PCs) that best separate networks. We then tested whether sex-biased clusters show significant differences across these PCs. We identify three PCs of cluster architecture that are significantly different (Relative Importance or RI >70%) between sex-biased and non-sex-biased clusters (Figure 5).

**Figure 5:**
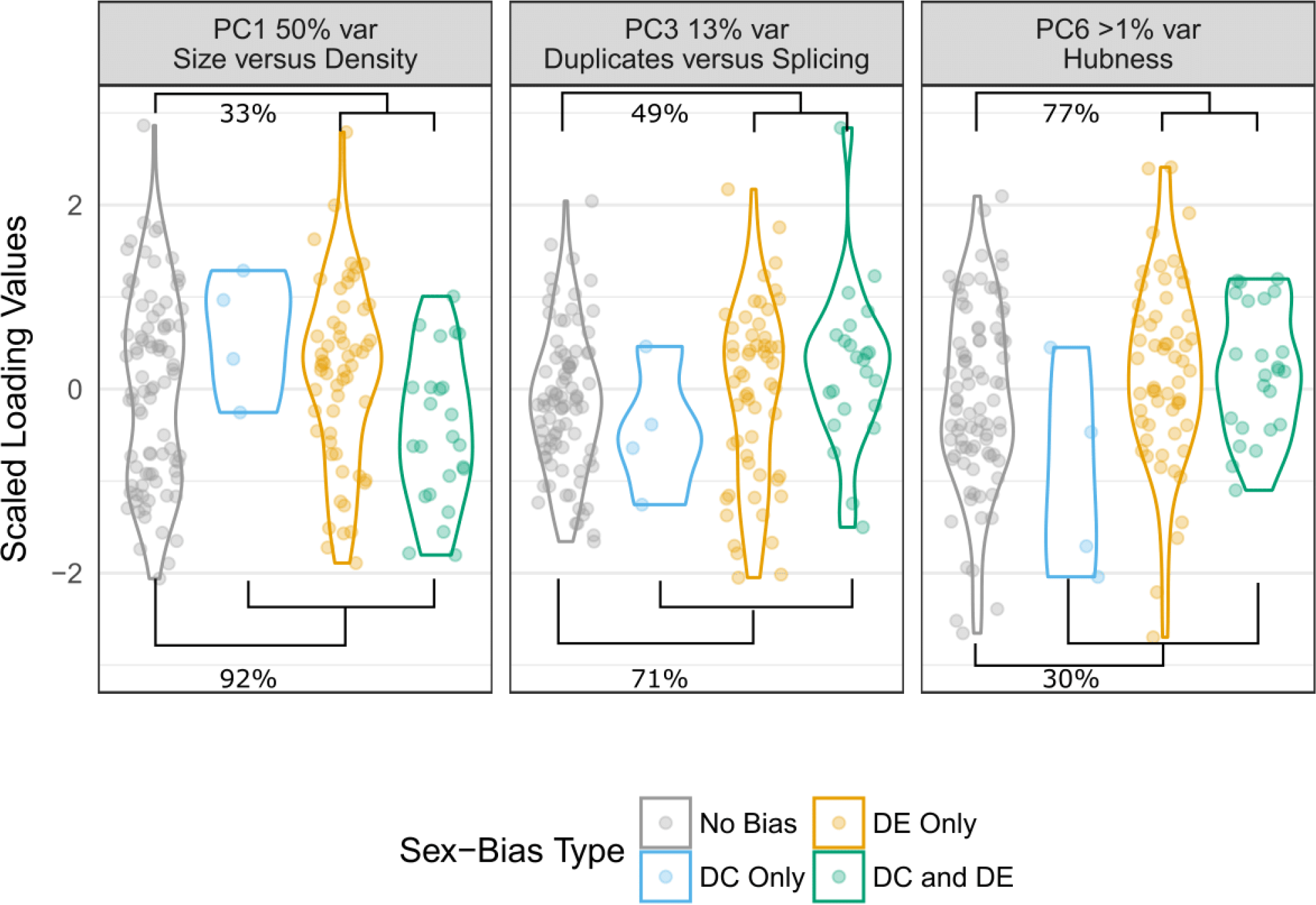
Topological differences between sex-biased and non-sex-biased clusters. Each panel shows one of the cluster parameters that differ between sex-biased and non-sex biased clusters. Percentages in each panel show the probability (RI) that a parameter differs between clusters with no sex-bias and clusters with differential expression (top of the figure) or differential correlation (bottom of the figure). The left panel discriminates between small clusters with many connections (negative values) and large clusters with few connections (positive values). The middle panel discriminates between clusters with many duplicated genes, but few genes with alternative splicing (positive values) and clusters with few duplicated genes but many genes with alternative splicing (negative values). The right panel discriminates between clusters with an uniform distribution of connection (negative values) and clusters with few genes with many connections and many genes with few connections (positive values).

The strongest association (RI 92%) is between differentially correlated clusters and PC 1, which is also the only factor with a significant ability to discriminate between clusters with sex-biased expression and clusters with sex-biased correlation (RI 93%). The lower scores of differentially correlated clusters on PC 1 indicate that they tend to have smaller sizes but higher density and a less centralized structure. High connection density has been shown to be related to greater network stability, since redundant connections buffer against mutations in the expression levels of individual genes (Levy and Siegal 2008; Rünneburger and Le Rouzic 2016). This allows the evolution of more stable phenotypes (Bergman and Siegal 2003) but tends to reduce their evolutionary potential (Wagner and Altenberg 1996). Accordingly, the higher density of differentially correlated clusters would predict lower evolutionary potential via network re-wiring in differentially correlated clusters compared to both differentially expressed and non-sex-biased clusters.

Differentially correlated clusters are also moderately associated (RI 71%) with PC 3, which is positively correlated with enrichment in duplicated genes and negatively correlated with enrichment for splicing nodes. This finding is in accordance with theories and empirical observations on how gene duplication can solve sexual conflict at the gene level (Baker et al. 2012; Gallach and Betrán 2011; Parsch and Ellegren 2013; Wyman, Cutter, and Rowe 2012; Bonduriansky and Chenoweth 2009), but are supported only for differentially correlated clusters. Taken together with the putatively low potential of evolution by re-wiring, the enrichment in duplicates could indicate that these clusters evolve primarily by adding new genes to the existing network rather than by re-arranging regulatory interactions between existing genes.

While PC 7 explains less than 1% of between-cluster variance, it is the only PC significantly associated with differentially expressed clusters (RI 77%). PC 7 is positively correlated with cluster centralization and negatively correlated with median clustering coefficient. The high PC 7 scores of differentially expressed clusters indicate a more hierarchical structure, with a stronger divide between hyper-connected hubs and peripheral worker nodes. This result suggests that differentially expressed clusters have a more ‘star’ shaped topology, in which most nodes are connected to hubs but not among each other. Thus, while differentially expressed clusters have average distribution of densities, the reduced amount of interactions between any given pair of genes could still be promoting rapid turnover of regulatory interactions (Wagner and Altenberg 1996; Mayer and Hansen 2017).

### Sex-Biased Clusters Integrate New Genes in Regulatory Positions

To validate whether sex-biased clusters show more rapid evolution compared to non-sex-biased ones, we initially compared the times of phylogenetic origin (stratum) of genes in both categories using data from Sackton, Werren, and Clark (2013). Reference species include non-chalcid hymenopterans (7 ants and 5 bees), insects, arthropods, and other metazoans (Figure 6, and see materials and methods for full species list). Compared with non-sex-biased clusters, sex-biased clusters show a greater proportion of genes that originate at the youngest taxonomic level (here chalcid). Among sex-biased clusters, those with differential correlation (DC) show a slightly greater proportion of chalcid-stratum genes than clusters with differential expression (DE). Compared to clusters that show only DE, DC clusters appear depleted of genes from more ancient phylogenetic origins (or strata), such as Hymenoptera, Insecta and Metazoa. This may indicate that clusters that show sex biased co-regulation are composed of genes that evolve relatively quickly, and therefore appear to be “younger” due to difficulties of identifying homology in deeper phylogenetic strata.

**Figure 6:**
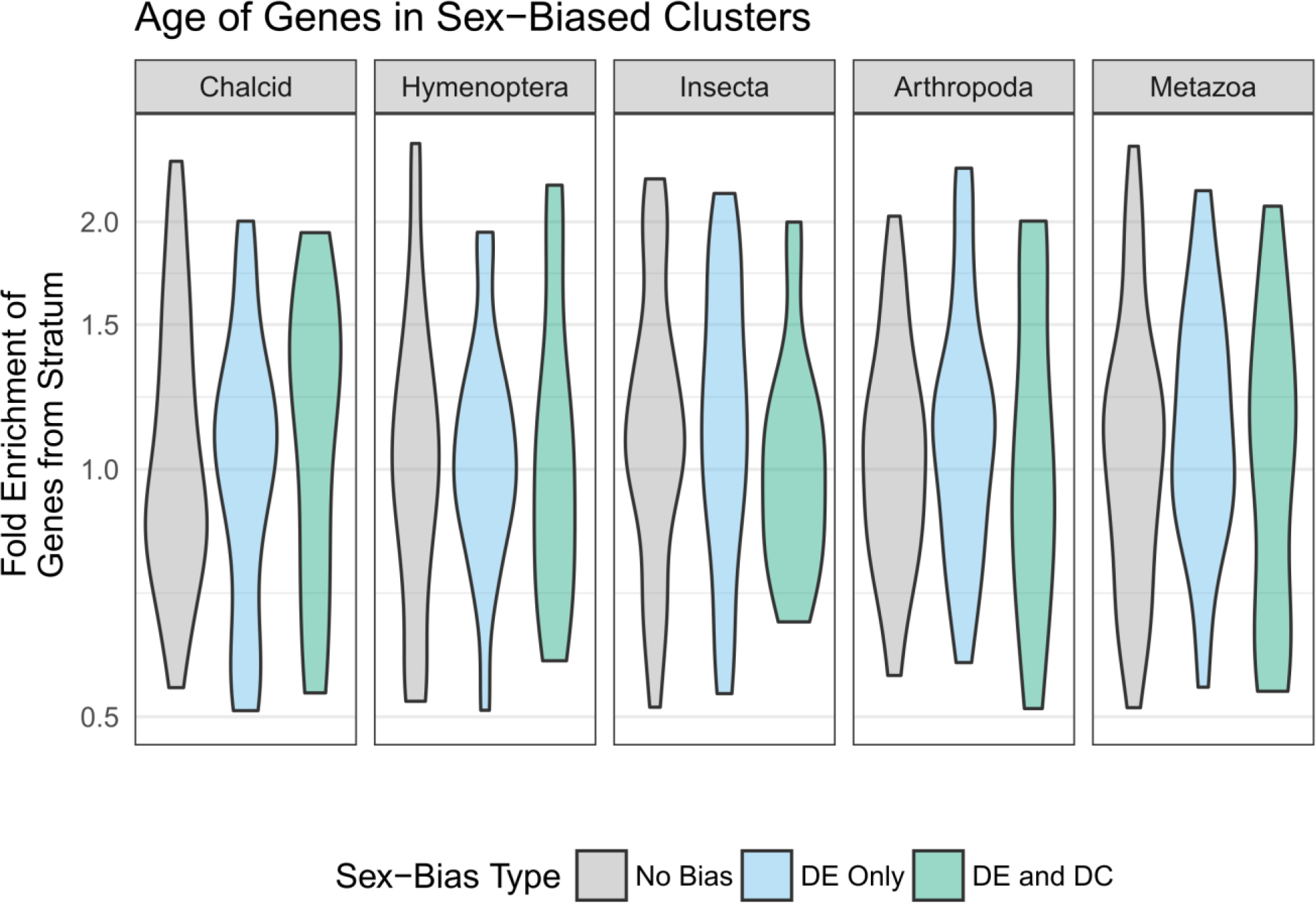
Age of genes in sex-biased clusters. Each panel shows the proportion of genes of a specific age in clusters with different types of sex-bias, compared to genome-wide averages. Gene age is measured as the highest taxonomic level required to describe all organisms that have homologs for the gene. Positive y values indicate that the clusters contain more genes of a specific age compared to genome-wide average (enrichment), measured in relative fold-change. For instance, a value of 1.5 indicates that the cluster contains 50% more genes of that age than expected from genome-wide average.

We next addressed whether new genes in sex-biased clusters are more likely to be in regulatory positions than new genes in non-sex-biased clusters. We compared the impact of gene age (phylogenetic stratum) with two main network properties: within-cluster connection density and hub scores. Connection density measures the number of interactions between the gene of interest and other members of its cluster. Hub scores estimate the regulatory potential of the gene of interest within its cluster (see Methods section and Figure 8 for details). We tested whether either of these two parameters is associated with the gene’s age (stratum), and whether the gene is part of a sex-biased cluster, as well as the interaction between gene age and sex-bias (Figure 7 and File S4). Since DE and DC clusters show different topologies (see previous section), we analyzed them as separate classes of sex-bias. We chose to also include cluster size as a predictor, since it strongly correlates with several features of overall cluster topology (see previous section and Figure 9), thereby controlling for potential interference from cluster-wide effects.

**Figure 7:**
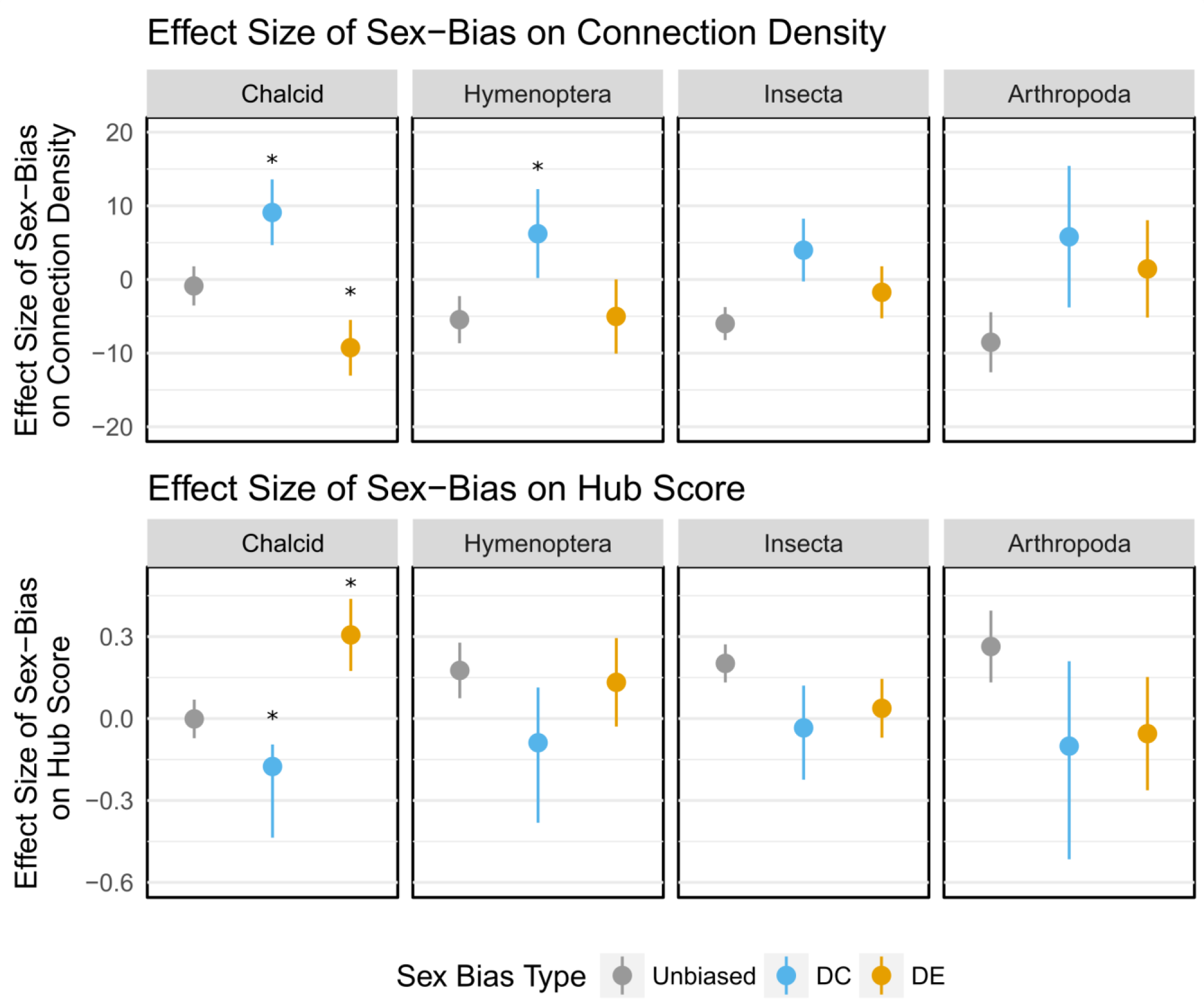
Topological properties of sex-biased transcripts of different ages. Each panel shows the average difference in connection density (top row) and hub score (bottom row) of transcripts from a specific age class compared to transcriptome-wide averages, with 95% confidence intervals. Colours show which transcripts belong to clusters with sex-biased expression, sex-biased correlation or no sex-bias. Contrasts marked with an asterisk indicate that transcripts in clusters with sex-biased expression or correlation have different connection density (top row) or hub score (bottom row) than transcripts in unbiased clusters for that age class. All contrasts are relative to the metazoan age class. See Supplementary File S4 for all coefficients.

We find that cluster size has significant effects on both density and hub scores (RI 100%), but in both cases, effect sizes are an order of magnitude smaller than those of age and sex-bias. Gene age is strongly associated with both connection density and hub scores, as indicated by its high relative importance in both models (RI 100%). Overall, genes from intermediate strata (Arthropoda, Insecta, and Hymenoptera) show lower connection densities and higher hub scores than younger genes, suggesting that genes from these strata are most frequently located in positions with high regulatory potential within their clusters. This pattern is reversed for genes assigned to the oldest stratum (Metazoan), which show high connection densities paired with low hub scores. This latter configuration is characteristic of interactor nodes (see Methods), which function by closely interacting with each other, rather than by directly regulating downstream elements. Accordingly, Metazoan stratum nodes are enriched by protein complexes (GO:0043234, q-value ~3.1 *e*^−96^) such as flagellar proteins in clusters ‘tomato’ and ‘skyblue3’, and spindle formation in cluster ‘thistle’.

On average, genes within sex-biased clusters show small but significant (RI 100%) increases in both density and hub scores compared to genes in non-sex-biased clusters. By contrast, genes within DE clusters show a marked decrease (RI 97%) in their connection density, consistent with their overall sparsity. Genes in DC and DE clusters also show a different relationship between age and topological properties compared to genes in non-sex-biased clusters, but the direction and magnitude of these interactions differs between DC and DE.

Genes in DC clusters show further decrease of connection densities with increasing gene age than non-sex-biased clusters (RI 100%) but maintain the overall pattern of increased hub scores with gene age as in unbiased clusters (RI 23%). The position of young genes in DC clusters is thus consistent with that of interactor nodes (see Methods), with low regulatory potential. By contrast, the relationship between gene age and topological properties is reversed for genes in DE clusters: younger genes show both decreased connection densities (RI 100%) and increased hub scores (RI 95%). This distribution indicates that younger genes in DE clusters conform to the expectation of hubs (see Methods), which bridge connections between otherwise independent group of genes and enable coordinated regulation.

To more precisely characterize the age of the young regulators in sex-biased clusters, we integrated phylostratigraphic assignments from additional parasitoid wasp species (Lindsey et al. 2018) in the superfamily Chalcidoidea. In addition to Nasonia (family Pteromalidae), the chalcids used here are those with published genome assemblies, and belong to different chalcid families, *Ceratosolen solmsi (Agaonidae fig wasps)*, *Copidosoma floridanum* (Encyrtidae), and *Trichogramma pretiosum*, in the more basal family Trichogrammatidae. The most recent common ancestor of these parasitoids is estimated to be approximately 99 MYA (Peters et al. 2018). These data allowed us to further separate genes from the hymenopteran stratum into genes that are shared among the chalcids (but not the other hymenopterans – Chalcid stratum) and those that are unique to *Nasonia* but not found in other chalcids (*Nasonia* stratum). The split between the *Nasonia* clade and its closest chalcid among our set (*Ceratosolen solmsi*) is approximately 71 MYA, followed by Copidosoma floridanum at ~81 MYA (Peters et al. 2018). Even after using this finer scale, only *Nasonia*-stratum genes show increased connectivity and hub scores in DE clusters relative to hymenoptera-stratum genes (Supplementary File S5), indicating that these features have evolved in the ~70-80 million year span that separates *Nasonia* (family Pteromalidae) and the most closely related Chalcid wasps included in our analyses (Martinson et al. 2017; Peters et al. 2018). While we did not detect the same pattern for DC clusters at this level of analysis, this is most likely due to the low number of genes with DC patterns that are assigned to the Chalcid stratum.

The different position of younger genes in differentially expressed and differentially correlated clusters is consistent with the global topological properties of these clusters. High density differentially correlated clusters show low capacity to evolve new regulators. By contrast, low density and high hierarchy differentially expressed clusters seem to allow the rapid integration of new genes with topological characteristics typical of regulators.

## DISCUSSION

Our assessment of sex-biased gene regulation across the development of *N. vitripennis* leads to several discoveries. We find relatively few differentially expressed (DE) genes early in development (early embryo, later embryo and larva). In contrast, 4 clusters show early sex-biased differential correlation (DC). These DC clusters contain another category of genes that can be important in early stage sex determination and differentiation, but which are normally overlooked in preference for genes showing highly biased differential expression (DE).

A dramatic increase of DE genes occurs in the pupal stage, when considerable sex differential development is occurring, and continues into adulthood. At the gene level, we observe a prevalence of sex-biased transcription relative to sex-biased splicing. Furthermore, genes are generally consistent in their sex bias during development. We find only a small number of genes that shift between male and female-bias across developmental stages, suggesting developmental constraints on sex-specific gene regulation. Most of the genes that switch between male and female bias may be attributed to differences in the timing rather than function between sexes (e.g. gonad maturation occurs earlier in males than females). We also identify several genomic regions enriched in male and female-biased genes, although their biological significance remains unclear.

At the cluster level, we report two distinct types of sex-biased clusters with specific temporal expression patterns and topological properties. Differentially correlated clusters show a surprising amount of cryptic sexual dimorphism in the earliest developmental stages. Differentially expressed clusters have a more hierarchical structure, with new or fast-evolving genes in key regulatory positions. Both DC and DE cluster genes provide reasonable candidates for future studies on sex-differentiation within the hymenopteran lineages.

### Prevalence of whole-gene transcriptional bias

Our analyses at the gene-level suggest that regulation of whole-gene transcriptional levels may be the most frequent molecular process inducing transcriptome-wide differentiation between the sexes. We find far more loci with evidence of sex-biased gene expression than sex-biased splicing. More importantly, the majority of genes with sex-biased transcription do not show sex-biased splicing, whereas most genes with sex-biased splicing also show sex-biased transcription. This inequality suggests that transcriptional regulation might be the prevalent method of solving within-locus sexual conflict, while sex-biased splicing may evolve less readily, or at a later stage in the resolution of sexual conflict. This finding is consistent with studies in *Drosophila* development (Brown et al. 2014), which show that the majority of splice variation is observed either between tissues or between stages, and that the few consistently sex-specific splice variants in adults can be attributed to sex-specific tissues.

Nonetheless, we describe 1,294 (~10% of 14,149) genes that show sex-biased splicing and lack sex-biased transcription. This finding doubles the reported frequency in adult *Drosophila* (Brown et al. 2014), yet is consistent with earlier *Drosophila* estimates from studies aiming at the specific detection of sex-biased alternative splicing (Telonis-Scott et al. 2008; Hartmann et al. 2011). It is also noteworthy that Brown et al. (2014) measured transcript expression via RNAseq technologies whereas both earlier *Drosophila* studies and our study relied on microarrays. As such, more molecular data is required to validate our findings on the scope of sex-biased alternative splicing.

Despite the abundance of sex-biased transcripts, only one gene (feminizer = *transformer*) shows consistent sex-bias across all developmental stages, while the majority of sex-bias is observed in either the pupal or adult stages. A study comparing pre-gonad imaginal discs in larvae and pupae to adult gonads in Drosophila found that 50-60% of sex-biased genes retain their expression-bias across larval, pre-pupal and adult stages (Perry, Harrison, and Mank 2014). However, that study is not comparable to ours since we compare whole body expression from early embryo to adult stages. Our estimates are closer to those observed in a study of chickens (Mank et al. 2010), which includes embryonic stages but again is focused on gene expression in gonads as opposed to the whole body study conducted here. The broad pattern is also potentially in accord with observations for caste-specific genes in ants (Ometto et al. 2011).

We find three main genomic regions enriched by sex-biased genes. A possible causal explanation of genomic co-localization for sex-biased genes is that regions of lower recombination promote co-adapted sex biased gene evolution. Co-expression of closely related genes may arise as a side-effect of tandem duplication, which can generate a large number of neighboring genes that share expression pattern because of identity by descent. This seems to be the case for the male-biased linkage group 4.1, in which a series of tandem duplications for male-biased “venom allergen” proteins (orthologous group EOG8W9MM2) is present. Since male *Nasonia* do not produce venom, we find it more likely that these proteins might be a component of male seminal fluids, a function that has indeed been reported for their Drosophila homologs (Kovalick and Griffin 2005) and would be consistent with the presence of secretory domains.

### Heterochrony in Gametogenesis Drives Developmental Sex-Bias Shifts

Only 3 clusters show shifts between male and female-bias at different developmental stages, suggesting that developmental sexual conflict significantly constrains sex-biased gene expression. However, these few clusters include 324 genes (1.1% of the whole network) and all three clusters shift from male-bias in pupae to female-bias in adults. These directional shifts in sex-bias may be explained by the timing of *Nasonia*’s gametogenesis. *Nasonia* spermatogenesis peaks during pupation, while its oogenesis occurs primarily during the adult stage (Whiting 1968). As such, we expect that genes required for gametogenesis will be expressed during pupation in males and during adulthood in females. The change in sex bias from male to female after pupation could thus be caused by gametogenesis genes that are not yet expressed in females during pupation, or have already been “switched off” in adult males. Genes in these male to female clusters are thus most likely performing the same functions at different times (heterochrony) rather than different functions in each sex.

While we identify several genes with female to male shifts in sex-bias (see Supplementary File S1 for details), this expression pattern is not represented in any cluster, suggesting that these genes may be outliers rather than members of functional sub-networks. These patterns contrast with those found for some other insects. By comparison, *Drosophila* shows sex-bias shifts in development in 4.9% of autosomal genes and 2.9% of X-linked genes (Perry, Harrison, and Mank 2014), while a study on *Bombyx mori* reports them for 54% of its transcriptome (Zhao et al. 2011). In both cases, the observed shifts are primarily (*Drosophila*) or exclusively (*Bombyx*) from female-bias in early stages to male-bias in the latter ones.

We cannot rule out that the increase of female expression in adults may be due to the greater proportional mass of gonads present in adult females compared to males. Characterization of clusters that show female-biased DE and male-biased DC reveals enrichment in testes-related processes, suggesting that (at least in these cases) tissue-bias in adults is strong enough to reverse measured gene expression-bias. To the best of our knowledge, only one study to date separated gonads from the carcasses in males and female *Nasonia* before RNA sequencing (Tennessen et al. 2014). While their findings are similar to those obtained by whole organism sequencing in Wang, Werren, and Clark 2015, the study focused on genes with at least 100-fold expression differences – effectively pre-selecting only for sex-specific genes rather than sex-biased ones.

### Sex-Bias in Early Development

We identified several cryptic early regulatory events using complementary analyses based on both differential expression (DE) and correlation (DC). Embryonic stages in particular show little differentiation between sexes when relying exclusively on differential expression. Yet several hidden co-regulatory events are revealed by the differential correlation analysis. For instance, only one cluster containing 79 genes is differentially expressed in early female embryos, compared to three differentially correlated clusters containing a total of 373 genes. Differential cluster expression identifies no male-biased genes in early embryos, whereas differential cluster integration reveals 75 male-biased genes. This result suggests that small proportional changes in the expression of multiple transcripts may play a previously unrecognized role in early sexual differentiation.

Among the clusters with female-biased correlation in the early stages, two are also differentially under-expressed in adult females relative to males. This pattern is potentially indicative of paternal imprinting mechanisms that influence early development. However, sperm is already mature in adult males, and therefore the opportunity for genomic imprinting is passed. It is possible that effects are mediated through seminal fluids. An alternative is that the clusters reflect induction of developmental pathways in early embryos derived from fertilized eggs, which are destined to become females. In *Nasonia*, male appears to be the default sex, with feedback between maternal derived and zygotic *doublesex* being required for induction of female development (Verhulst et al 2010; 2013). Under-expression of these genes in the adult female (e.g. ovaries) could reflect a male default pattern, with induction of the female developmental cluster through *doublesex*. Therefore, the genes in these DC clusters may be important for this early sex differentiation.

Histone and histone-modification enzymes are enriched and occupy hub positions in the only early embryonic cluster that also shows sex-bias according to both our measures. Overexpression (DE) of histones in diploid females may be expected due to the greater amount of DNA in their cells but DC is caused by female-specific increase in expression correlation, which is independent from absolute expression. The presence of DC suggests that histones and their modification enzymes may be involved in sex-specific interactions in early embryogenesis. This result is especially interesting in light of the ongoing debate on *Nasonia*’s sex-determination mechanism. While there is now consensus on the need for a silencing mechanism of maternal Feminizer expression (Verhulst, van de Zande, and Beukeboom 2010; Verhulst et al. 2013), the specific mechanism has so far been elusive. Several papers aimed at investigating the role of DNA methylation have shown that genes subject to DNA methylation show less variation across evolutionary and developmental space (Park et al. 2011; Wang et al. 2013) and there is very limited evidence for sex-biased differential methylation in adults (Wang, Werren, and Clark 2015). Our study reinforces a lack of support for DNA methylation as a mechanism for sex-biased genome imprinting, suggesting instead the modification of specific histones as a possible alternative. Since the genome copy carried by sperm is bound by sperm-specific protamines (Tennessen et al. 2014) rather than histones, such a mechanism would provide a robust means of erasing only paternal imprinting without the need for divergent histone markings in adults. Histone-mediated *Nasonia*-specific control of sex-determination would also be consistent with our earlier finding that histone gene families are specifically expanded in Nasonia relative to ants and bees (Rago et al. 2016), and that some of those members are indeed part of the clusters with early sex-bias.

### Network Structure of Sex-Biased Clusters

Sex-biased clusters show high proportions of *Chalcid*-stratum genes (Figure 6), which occupy different positions within their networks (Figure 7). In differentially correlated clusters, *Chalcid*-stratum genes are highly connected but have low hub scores, indicative of low regulatory potential. This result is consistent with the hypothesis that dense regulatory networks would be constrained in the evolution of new regulators due to the higher proportion of pleiotropic interactions between their components (Pavlicev and Hansen 2011, but see Mayer and Hansen 2017), yet is restricted only to clusters which show sex-biased correlations rather than being present across the transcriptome.

Differentially expressed clusters are more hierarchically organized, as measured by their lower density and higher centralization. One of our initial hypotheses is that both sparsity (few connections) and hierarchy (strong division between regulators and workers) may facilitate the emergence of new regulators. The younger (or more rapidly evolving) *Chalcid*-stratum genes in differentially expressed clusters are sparsely connected and show the highest hub-scores. Their preferential position between groups of otherwise unconnected nodes is characteristic of gene regulators and reveals a propensity of differentially expressed clusters to incorporate new genes in control positions. While a sparser network would increase the odds of a new node to become a regulator, the prevalence of *Chalcid*-stratum nodes in hub positions remains surprising when compared to equally sparse non-sex-biased clusters, whose hubs are instead occupied by nodes from the Arthropoda and Insecta strata. This result may suggest rapid evolution of sex-bias networks relative to more stable non-sex-biased networks.

The method of phylostratigraphic dating can generate biases (Moyers and Zhang 2014), particularly when attempting to detect deep matches for short or rapidly evolving genes whose sequence similarity rapidly degenerates below homology criteria. Considering that sex-biased genes have indeed often been observed to have faster evolutionary rates (Wang, Werren, and Clark 2015), it is likely that a portion of *Chalcid*-stratum genes will consist of rapidly diverging genes from older strata. Depending on the extent of phylostratigraphic-bias, we can interpret these findings in two ways. Either new genes are indeed more readily integrated in key regulatory positions within differentially expressed networks (low phylostratigraphic-bias scenario) or genes in key positions of sex-biased networks in the *Nasonia* clade have rapidly mutated beyond homology criteria (high phylostratigraphic-bias scenario). Both scenarios imply that genes in sex-biased clusters show significant evolutionary differences compared to non-biased ones, and that those differences are closely related to the genes’ positions within the regulatory network. Rapid integration of novel genes into regulatory positions of sex-specific networks has already been documented multiple times in *Drosophila* for mechanisms as diverse as male fertility (Ding et al. 2010; Dai et al. 2008) and courtship specificity (Dai et al. 2008), whereas over 75% of the caste-biased genes in the social wasp *Polistes canadensis* lack homology outside of the species (Ferreira et al. 2013).

The pattern of relatively higher turnover of regulators acquisition in differentially expressed clusters that we observe is consistent with Developmental Systems Drift (True and Haag 2001; Haag 2014) – an evolutionary model that allows for the change of the underlying regulatory pathways *via* stochastic drift while preserving the phenotype through the repeated emergence and loss of redundant regulators. A similar pattern is already observed for the primary regulators of sex-determinations across Insecta and Hymenoptera (Verhulst, van de Zande, and Beukeboom 2010; Koch et al. 2014) and could be indicative of a general feature of sexual development. It is also possible that ongoing sexual antagonistic selection (i.e., optimal expression and correlation in one sex is suboptimal in the other), and genetic conflict between differentially heritable elements influencing sex determination, creates a motor for turnover in sex-biased gene network organization (Werren and Beukeboom 1998). With rapid rates of molecular evolution and a strong constraint for retaining two functional phenotypes, sexually dimorphic development might indeed be the optimal scenario for the prevalence of DSD.

## CONCLUSION

Here we present a rare investigation of sex biased gene expression and co-expression from early development (embryo) to adulthood in an insect. *Nasonia*’s sexual development reveals numerous interesting properties about the evolution of sexual dimorphism and identifies sets of candidate genes for early and stage-specific sex differentiation that provide further understanding on the evolution of sex-determination in Hymenoptera. We also make a first detailed comparison of the interplay between transcription and splicing over *Nasonia’s* sexual development, assessing the prevalence of transcription and noting instances of splicing that most likely mediate sexual conflict. Our analyses of early developmental expression reveal that differentially correlated sets of transcripts could play a previously unrecognized role in the onset of sexual differentiation and possibly sex-determination itself. Despite the lack of genetic sex-determination, we find at least three genomic regions enriched in sex-biased clusters. Several scenarios can explain their presence, spanning from selective advantage of their co-inheritance to non-adaptive linkage hitchhiking. Discriminating between these options will require modelling that integrates knowledge about *Nasonia*’s genome with its ecology and taxonomy.

*Nasonia*’s sex-bias is strongly developmentally restricted, with few transcripts showing sex-bias in multiple stages. While several genes show male to female sex-bias changes between stages, they remain mostly confined to meiosis-related genes or contrasts between pre- and post-pupation stages. The observation that the same genes tend towards the same sex-bias direction across development supports the presence of strong constraint in sex-bias across development. The prevalence of stage-specific sex-bias, and the fact that transcripts that shift in sex-bias direction do so mainly during pupation, underscores the importance of treating different life-stages as factors of interest to correctly understand gene expression evolution, and reflects developmental timing differences between the sexes and between their homologous structures.

Finally, our characterization of two main classes of sex-biased clusters via network analyses provides a better understanding of the role of novel genes within co-regulated clusters. While all sex-biased clusters showed enrichment for novel genes, we find them to occupy fundamentally different positions in their networks, acting as potential regulators only in differentially expressed clusters. This finding provides an empirical confirmation for hypotheses on how sparsity and hierarchy can facilitate the evolution of regulatory structures (Kouvaris et al. 2017; Mengistu et al. 2016). The observation that novel genes can be incorporated into pivotal regulatory positions in sex-biased clusters poses a challenge to the evo-devo assumption that regulators are conserved over time, supporting instead a model of phenotypic stasis and regulatory reshaping that characterizes developmental system drift (True and Haag 2001; Haag 2014), or alternatively, accelerated evolution of sex-biased gene networks due to sexual conflict.

## Materials and Methods

### Biological Materials and Data Collection

The data for this study consisted of a developmental time series of transcriptomes occurring in whole animals, both for male and female jewel wasps (*Nasonia vitripennis*). Our experiment was conducted using a highly-inbred strain called AsymCX (Werren et al. 2010). Inbreeding occurs routinely in *Nasonia* without the deleterious effects often found in outbreeding diploid species, primarily due to purging of mutational load in the haploid sex (Werren 1993) and provides a reduced level variation due to genetic differences between individuals.

We sampled animals at five distinct developmental stages: early embryo (0-10 hr old), late embryo (18-30 hr old), 1^st^ instar larvae (~51 hr old), yellow pupa stage (~14 days) and sexually mature virgin adults. Specifically, samples taken during the first embryonic stage were of single zygotes up to late blastoderms, prior to gastrulation. The samples taken during the late embryonic stage were of embryos that exited gastrulation and include segmentation, organogenesis and the remaining pre-hatching development (for reference timings, see Bull 1982).

Each of these developmental stages was sampled in triplicate for each sex, for a total of 30 samples to profile. Because of the different number of cells at different stages, different numbers of sampled individuals were pooled for each biological replicate as follows: 300-900 individuals for early embryos, 140-500 for late embryos, 245-520 for 1^st^ instar larvae, 20 for pupae and adults.

Pupae and adults were produced by mated females and sexed by visual examination prior to extraction and expression profiling. Since sexing by visual examination is impossible before the pupal stage, male embryonic and larval samples were collected from virgin females, which produce only males. Female embryonic and larval samples were collected from mated females, which produced ~83% female offspring.

Expression values were measured via dual-channel whole-genome tiling path microarrays using custom NimbleGen high-density 2 (HD2) arrays (Lopez and Colbourne 2011), consisting of 8.4 million probes with a 50-60 nt length spanning the *Nasonia* genome at 33 bp intervals, as well as 27,000 Markov probes which are absent from the genome for noise detection (see below). Further details on animal breeding, RNA extraction and microarray processing are available in the supplementary materials of (Werren et al. 2010).

### Data Pre-Processing

Individual microarray probes were assigned to exons according to the latest release of the Official Nasonia Gene Set (OGS2.0) (Rago et al. 2016). Expression for each exon was measured as the log2 ratio of the 99th quantile of the random Markov probes on their arrays. A sensible expression cut-off was set by examining the distribution of exon expression values across the whole experiment. Based on this assessment, all values below the 66th expression percentile were collapsed to zero to avoid spurious signal from random noise variation among non-expressed exons. We also retained only exons that showed expression above this signal threshold in at least two out of three replicates for at least one biological condition were retained for analysis.

Therefore, of the 126,213 annotated exons and 23,149 annotated genes from OGS2.0, only 67,409 exons and 14,151 genes were retained for investigating their transcriptional profiles after taking these quality assurance measures. Two more genes were filtered out as the result of quality control downstream in the pipeline: gene NASVI2EG009090 was removed during splicing detection (see below) since it showed low variation in expression across samples. Gene NASVI2EG021272 was removed before result interpretation because of its low annotation quality in OGS2.0. As such, the final dataset presented includes a total of 14,149 genes.

### Splicing Detection Identifying Transcription-Nodes and Splicing-Nodes

We applied the FESTA algorithm (Rago and Colbourne 2018) to disentangle transcription and splicing signals. Briefly, FESTA allows detection of alternative splicing based on experiment-specific exon expression data. FESTA disentangles alternative splicing signal (creating **‘splicing-nodes’**) from whole-gene transcription (**‘transcription-nodes’**) by first identifying constitutive exons that are present across all isoforms, then by measuring the relative frequency of facultative exons. We define facultative exons here as exons which are present in only some of the transcripts produced by each gene across the whole experiment. We choose to convert the expression values of splicing nodes to splicing ratios, which are normalized to the total expression value of the main gene within the same sample. Splicing ratios represent the proportion of all transcripts produced by a single gene which include the exons of interest. This allows splicing nodes to be represented as continuous proportions in a 0-1 range and removes the correlation between overall gene expression and expression of isoforms containing the exons of interest.

The final dataset contains a total of 36,505 nodes, 14,149 of which are transcription nodes and 22,356 of which are transcription nodes. Since each node is representative of a putative transcript, we refer to them using the term node and transcript interchangeably.

We subsequently collapsed all nodes with reciprocal correlation values greater than 95% into Constitutively Correlated Regulatory Events (CCREs) using the ‘collapseRows’ function from the WGCNA package implemented in R (Langfelder and Horvath 2008). This step helped to reduce the dimensionality of our dataset by representing sets of nodes with almost identical expression patterns as single units. Each CCRE thus represents the expression pattern of a set of transcripts which behave identically to each other within our experiment. This approach is conceptually similar to that of Constitutively Co-expressed Links (CCELs) in (Hsu, Juan, and Huang 2015). We chose to represent each CCRE using expression scores of the node with the most correlation-based connections to other nodes in the same CCRE, because this value is the most representative of the average behavior of other CCRE members. In the special case were CCREs contained only two nodes, we chose the highest mean expression value. A total of 15,792 nodes was grouped into 3,699 CCREs. As such, the final dataset features a total of 24,412 features of which 8,720 (36%) are transcription nodes, 11,993 (49%) are splicing nodes and 3,699 (15%) are CCREs. The complete annotation of all transcripts assigned to each CCRE is available in Supplementary File S3.

### Network Construction

We constructed an undirected weighted interaction network of transcription-nodes and splicing-nodes using the R package WGCNA (Langfelder and Horvath 2008). WGCNA infers between-node links based on power-transformed robust correlation scores. Since WGCNA does not require the input of pre-defined pathways or functional classes, it is ideally suited for the analysis of expression data from genomes with sparse functional gene annotations. WGCNA is also able to rapidly calculate large networks, which is a key feature for enabling permutation-based approaches to monitor differential correlations (see below).

Finally, we power-transformed pairwise correlation scores. This transformation increases the difference between weak and strong links in our network, with the effect of increasing overall specificity in detecting connected sub-networks (Zhang and Horvath 2005). Most natural network studies show a power-distribution of connectivity across nodes (Jeong et al. 2001; A. Wagner and Fell 2001; Barabási and Oltvai 2004), with few highly connected nodes and many lowly connected ones, also called a scale-free degree distribution. Based on this “scale-free topology criterion” (Zhang and Horvath 2005), we selected the lowest power that generated a scale-free correlation network.

### Network Topology Measurements

We measured two main network topology parameters for each node: connection densities and hub-scores.

**Connection density** is defined as the number of observed connections per node, normalized by the theoretical maximum possible number of connections. In an undirected weighted network, this parameter can be calculated with the formula 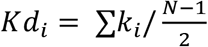 where ∑*k*_*i*_ is the sum of the weights for all connections to node *i* and 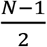 is the maximum number of links in an undirected network of size *N*. Connection density quantifies the relative importance of a node as a measure of its direct connections to other nodes within the same network (i.e. **Figure 8**, nodes A and C), and is useful to estimate the number of interactions among individual genes.

However, connection density does not account for the different regulatory potential of different connections. For instance, nodes A and B (**Figure 8**) have the same connection density. However, removing the connections from node A divides the network in two while removing the connections from node B causes no splitting in sub-networks, since its neighbors are already connected with each other.

**Figure 8:**
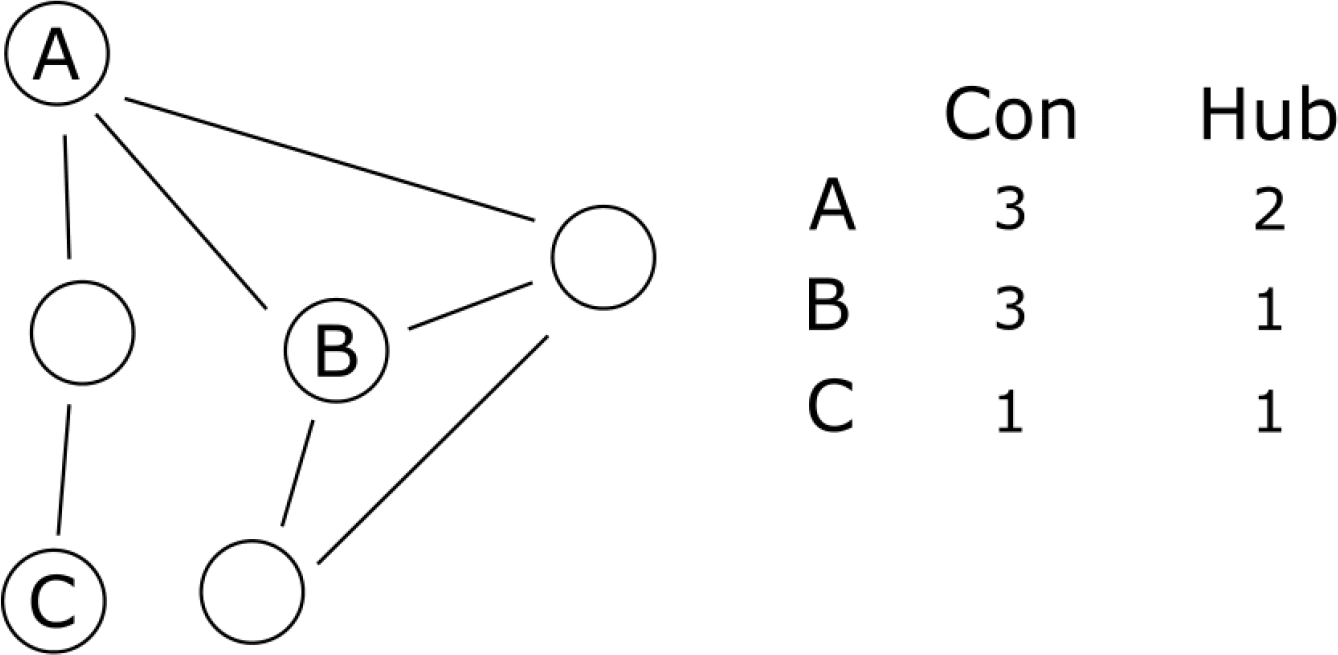
Topological properties of regulator, interactor and marginal nodes. Nodes represent individual transcripts; connections represent correlations in expression values. Node A (regulator) has high connection density and high hub score since it connects several nodes which are otherwise not connected. Node B (interactor) has high connection density and low hub score, since it only connects nodes that are already connected. Node C (marginal) has low connection density and low hub score, since it connects to few nodes that are already connected with others.

To account for this topological property, we calculated **hub scores**, which measure the non-redundant connections provided by each node. We calculated hub scores using the formula 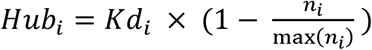 where *Kd*_*i*_ represents the connection density of node *i* and 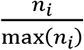represents the clustering coefficient of node *i*, or the observed connectivity between nodes connected to node *i* divided by their maximum possible connectivity with each other. Nodes with high hub scores have a high number of connections to nodes that are otherwise unconnected among themselves and are likely to be involved in the coordination of multiple processes. Since we calculate hub scores by penalizing connection densities, a node’s hub scores cannot be higher than its connection density.

We use these two topological parameters to define three main classes of nodes in our network: **Marginal nodes** have low density and low hub scores (**Figure 8**, node C), and are unlikely to be involved in regulation. **Regulator nodes** have high density and high hub scores (**Figure 8**, node A), and are likely to be key regulators. Finally, **interactor nodes** have high density and low hub scores (**Figure 8**, node B), and form potentially functional stable associations with other nodes but are unlikely to be regulators.

To focus our investigation on the structure of local regulatory sub-networks across the sampled developmental stages, we calculated both scores by considering nodes within the same transcriptional cluster. We computed the within-cluster network statistics for each node using the ‘fundamental Network Concepts’ function from the WGCNA package, as well as weighted betweeness using the tnet package (Opsahl 2009).

### Differential Expression of Nodes and Clusters

We detected differential expression (DE) of individual nodes using generalized linear models (GLMs) as implemented in the LIMMA package (Smyth 2005). We used the formula

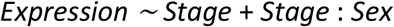

which accounts for stage-specific differences in gene expression via the factor *Stage* and considers sex only as a second-order interaction term with stage-specific expression changes (*Stage*: *Sex*).

To calculate cluster-level DE, we applied the same linear models to the first principal component (module eigengene) of each cluster. We performed multiple-hypothesis correction on both node and cluster-level DE results by converting the individual p-values to local False Discovery Rates (lFDR), which represent the individual probability of each hypothesis to be a false positive. We calculated lFDR using the R package fdrtool (Strimmer 2008) on the p-values generated by LIMMA. All contrasts with a lFDR lower than 5% were considered significant. We classified each gene as sex-biased if at least one of its transcription- or splicing-nodes was differentially expressed (DE) between the sexes in at least one developmental stage. Full results for node-level analyses are included in Supplementary File S1.

### Linkage Clusters Enriched by Sex-Biased Loci

We next investigated whether sex-biased genes were enriching known linkage intervals, which would be indicative of sex-determining regions (SDRs) within the *Nasonia* genome. Since each node is DE tested independently at each stage, it is possible for a single gene to be both male and female-biased at different stages. Likewise, different transcription and splicing nodes from the same gene can show bias in either sex. Genes that fall in either category are unlikely to be subject to sex-specific selection, so we excluded them from linkage group enrichment analyses. We mapped all genes in our network to the linkage map published in (Desjardins et al. 2013), then tested each individual linkage group for enrichment in male or female-biased genes via one-tailed Fisher’s exact test, compared to the overall proportions of male and female-biased genes across all other linkage groups. This process generated two p-values per linkage group: one for female bias enrichment and one for male-bias enrichment. Finally, we applied FDR correction to the p-values using the package fdrtool, and reported all clusters with a lFDR score lower than 5%.

### Differential Correlation Analyses

In contrast to detecting differential expression of nodes based on significant changes in the relative abundance of transcripts, the detection of differential co-expression classifies groups of genes as biologically interesting based on a differential increase or decrease in their correlations across the conditions of interest (Figure 1). There are two distinct methods used to analyze differential co-expression: **‘untargeted’** methods identify changes in transcript-transcript interactions (Tesson, Breitling, and Jansen 2010; Ma et al. 2011; Hsu, Juan, and Huang 2015; Liu et al. 2016), and **‘targeted’** methods measure correlation changes in pre-defined groups of transcripts (Yang et al. 2013; Cao et al. 2014). To allow direct comparisons between differential correlations and differential expression data, we developed a targeted method that is applied to the co-expression clusters found via network reconstruction, a strategy is termed **‘semi-targeted’**.

The development of a semi-targeted strategy was necessary because most available methods are designed for two-sample tests, or to detect individual sample deviation from a pre-defined baseline (Yu et al. 2011; Walley et al. 2012; Liu et al. 2016), and are thus unable to account for multi-level and nested experimental designs. Moreover, untargeted methods are too computationally costly and underpowered for the comparisons of cluster-level DE.

Since our objective is to detect sex-specific gene co-regulation, we employed a sub-sampling strategy that removes all possible combinations of three samples within each stage, which equals the number of biological replicates for each stage by sex combination. This sub-sampling strategy retains a constant number of samples used for the generation of each sub-network, while systematically altering the proportion of samples from each sex at each developmental stage. For each sub-sampling iteration, the sub-networks were reconstructed using the R package WGCNA (Langfelder and Horvath 2008) using the same power transformation and node to cluster assignments as for the main network reconstruction (see **Network Construction** above). We then measured the within-cluster density of each cluster in every sub-sampled network. Since WGCNA-based cluster density is effectively a power-transformed measure of correlation between nodes in a cluster, we refer to its differential change as **‘differential correlation’** (DC).

Within-cluster density is a proportional measure that is distributed on a scale of 0 to 1, where 1 indicates that all possible connections between nodes are observed and where 0 indicates that no connections between nodes. The within-cluster density can therefore be analyzed using Generalized Linear Models (GLMs) with a gamma error distribution and logit link function. We fitted the following GLM to each cluster:

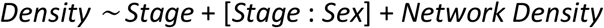

This approach allowed the detection of stage-specific sex-bias in cluster density [*Stage*: *Sex*] while controlling for stage-specific and whole-network increases in connectivity. To validate that the observed density bias is non-random, we fitted the same GLM to 1000 datasets generated by randomly permuting sex-labels. The p-values for the [*Stage*: *Sex*] interactions were then calculated for each cluster from the GLMs of both the permuted and observed datasets. We then estimated the significance threshold for each case of differential correlation by calculating the lFDR of observed [*Stage*: *Sex*] p-values compared with the distribution of p-values generated by the randomly permuted dataset. Finally, we corrected for multiple-hypothesis testing by calculating, for each cluster, the [*Stage*: *Sex*] lFDR score against all other cluster lFDR scores. All [*Stage*: *Sex*] interactions with a lFDR score lower than 10% were deemed significant, leading to an expectation of making less than 2 false discoveries.

### Multivariate Analysis of Network Parameters

Network and sub-network parameters display several non-trivial correlations (Horvath and Dong 2008; Dong and Horvath 2007). Consequentially, we observe strong non-independence between our parameters of interest (Figure 9). We therefore employed a Principal Component Analysis (PCA) to detect the latent independent components that affect network parameters. The factors included in the PCA analysis are: cluster size (number of nodes), density, centralization, heterogeneity and median cluster coefficient as defined in (Horvath and Dong 2008), as well as cluster diameter (the longest among shortest paths within the network). We also included the relative proportion of splicing nodes and the relative proportion of nodes arising from duplicated genes as factors in the PCA analysis. Both of these proportions were normalized by their respective network-wide abundances before PCA. All variables were also centered and scaled before PCA.

**Figure 9:**
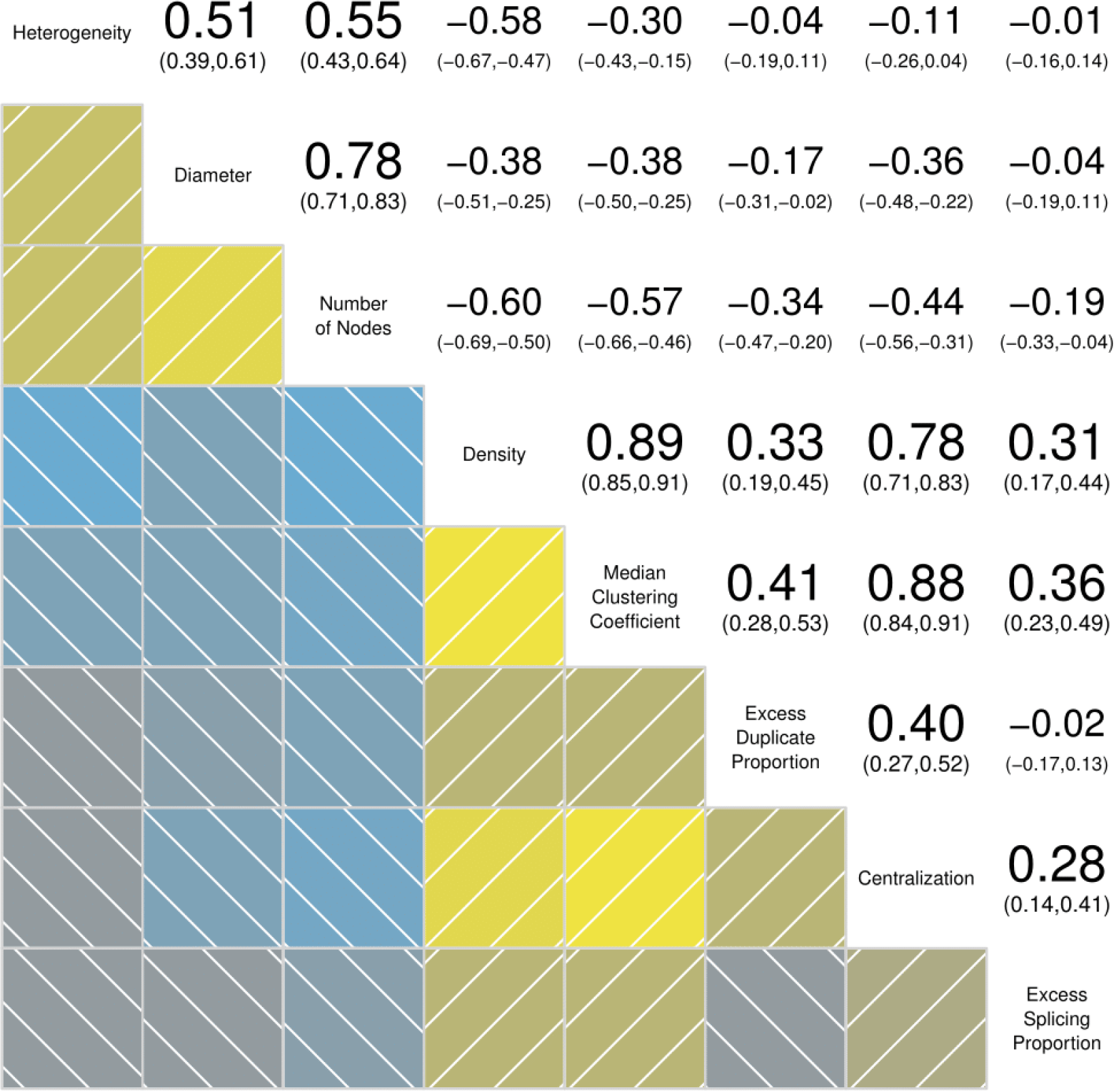
Correlations between topological network parameters. Yellow squares in the bottom left corner indicate positive correlations, blue ones negative. Lighter shades are more significant than darker ones. Numbers at the top right corner indicate the Pearson correlation score with confidence intervals in parentheses.

Each of the principal components (PCs) extracted by PCA represents a single linear combination of the factors, which maximizes the degree of variance between clusters, and minimizes the reciprocal correlation with other PCs. We determined the biological significance of each PC by comparing the relative contribution of each parameter to their score (as estimated by parameter loadings). Since our objective is to determine whether any of the latent variables can discriminate between the different classes of sex-biased clusters, we used binomial GLMs including all PCs as predictors. We thus fit three separate model sets, using the following dependent variables: differentially expressed cluster, differentially correlated cluster, clusters with both differential expression and correlation. We then computed model sets containing all possible combinations of factors for each of the three main models and estimated each factor’s probability of being included in the best model of its set (their **relative importance**, or **RI**) using the small sample-size corrected version of the Akaike information criterion (AICc) based rankings as implemented by the R package MuMIn, (Barton 2011). The results for differentially correlated clusters and clusters with both differential expression and differential correlation were identical (data not shown), most likely because only 4 clusters with differential correlations showed no differential expression in at least one stage. Due to this matching, we only report results for the model set targeting differentially correlated clusters.

To detect whether any PC differs significantly between differentially expressed and differentially correlated clusters, we fitted a fourth binomial model set including only clusters with either differential expression or differential correlation, using differential correlation as a dependent variable and the eight PCs as its predictor.

### Phylostratigraphic Analyses on Network Parameters

We used phylostratigraphic analysis to investigate the evolutionary age of genes within our networks (Domazet-Lošo and Tautz 2010) using annotations for *Nasonia* from (Sackton, Werren, and Clark 2013) for hymenoptera and higher levels. Briefly, each gene is assigned the phylogenetic stratum of its deepest detectable homolog. Two genes are considered homologs if they have a blastp hit whose alignment covers at least 50% of the shorter protein with at least 30% positives (see Sackton, Werren, and Clark 2013).

The hymenopteran species used are the ants (*Formicidae*) *Atta cephalotes, Acromyrmex echinatior, Camponotus floridanus, Harpegnathos saltator, Linepithema humile, Pogonomyrmex barbtus, Solenopsis invicta* and the bees (*Apidae*) *Apis florea, Apis mellifera, Bombus impatiens, Bombus terrestris, Megachile rotundata. Nasonia* is the only representative of the highly diverse and speciose Chalcioidea superfamily (J. M. Heraty et al. 2013; Peters et al. 2018). Therefore, in this analysis, we assigned Nasonia genes without orthologs in the other hymenopteras the “chalcid” stratum. Higher level stratum assignments were based on 11 additional insect, 2 additional arthropod, and 5 additional metazoa species.

We performed a second identical set of analyses using the subset of genes included in the hymenopteran and chalcid strata, and adding more species to increase the phylogenetic resolution within hymenoptera. We used data from Lindsey et al. (2018) for homology assignments to the chalcid species *Ceratosolen solmsi*, *Copidosoma floridanum*, and *Trichogramma pretiosum*, as well as the additional hymenopterans *Athalia rosae*, *Orussus abietinus*, and *Microplitis* demolitor. These additional data allowed separating genes that are found within chalcidoid wasps (matching with *Ceratosolen*, *Copidosoma* or *Trichogramma*), but not in the other Hymenoptera. Genes were assigned to the Chalcid stratum when an ortholog was found in one or more of the other chalcidoids, and to the Nasonia stratum when the Nasonia gene did not have any ortholog. Orthologs for this set were assigned using OrthoMCL with an e-value cutoff of 10e-10, percent match cutoff of 70 and inflation of 2.5 (Lindsey et al. 2018, Supplementary Information, section S2.2).

Since few *Nasonia* genes from our prior stratum analysis matched to species within the Apocrita stratum (*Apis mellifera* and *Micropolitis demolitor*), we merged it with the closest older stratum (Hymenoptera). *Nasonia* genes which did not have homologs with any species within the set were assigned to the *Nasonia* stratum. All stratum assignments are available in Supplementary File S1.

We used GLMs to test for the impact of the inferred phylostratigraphic age on each node’s within-cluster connection density and hub scores by fitting the following models to each of the datasets:

1. *Connection Density ~ Cluster Size* + *Stratum* + *DE* + *DC* + [*Stratum*: *DE*] + [*Stratum*: *DC*]
2. *Hub Score ~ Cluster Size* + *Stratum* + *DE* + *DC* + [*Stratum*: *DE*] + [*Stratum*: *DC*].

These models estimate the ability of taxonomic strata to predict connection density and hub scores both independently (term *Stratum*) and by interacting with two main sex-biased parameters (terms [*Stratum*: *DE*] and [*Stratum*: *DC*]), after controlling for variation in connection densities due to sex-bias parameters (terms *DE* and *DC*) and cluster size. Since connection densities and hub scores are expressed as 0-1 bound variables, we used a gamma error distribution and a logit link function. We subsequently fitted all possible nested models and produced model-averaged parameter estimates and RIs for each factors using AICc based rankings (as implemented in Barton 2011). Results from the Metazoan to hymenoptera stratum assignments are show in supplementary file S4, and results from the Hymenoptera to *Nasonia* stratum assignments in supplementary file S5.

### Gene Ontology, and Protein Family Enrichment Analyses

For the Gene Ontology (GO) and PFAM (Protein Family database) enrichment tests, we used the interface provided by Wasp Atlas, which returns FDR-corrected q-values for over-representation of GO and PFAM categories in the gene-set of interest by using one-tailed FDR corrected hypergeometric over representation tests (Davies and Tauber 2015). We choose a significance threshold of q < 0.01. The input files used for enrichment testing were either lists of genes (for linkage group enrichment) or transcription nodes (for transcriptional cluster enrichment).

## ACKNOWLEDGEMENTS

The research was supported by US National Institutes of Health grant to JHW (1R24GM084917 “Genetic and Genomic Tools for the Emerging Model Organism, *Nasonia*”) including a sub-award to JKC. Additional support was provided to JKC by the University of Birmingham, and to JHW by the US National Science Foundation (IOS-1456233) and Nathaniel & Helen Wisch Chair. Special thanks go to R. Edwards and J. Lopez for assistance with the experiments, plus D. Gilbert and J.-H. Choi for the microarray annotation.

## SUPPLEMENTARY FILES

File S1: Complete annotation of *Nasonia* transcripts described in the experiment. File S2: Complete annotation of *Nasonia* transcriptional clusters.

File S3: Annotation of transcripts comprising Constitutively Co-expressed Regulatory Events.

File S4: Coefficients and importance of phylostratigraphic age on network parameters, using strata from Metazoa to Chalcid.

File S5: Coefficients and importance of phylostratigraphic age on network parameters, using strata from Hymenoptera to *Nasonia.*

