## Supplementary Files 1-5 for "Sex biased expression and co-expression networks in development, using the hymenopteran *Nasonia vitripennis*": S1_Readme.txt

transcriptID		ID of the transcript, as detected via the FESTA algorithm
geneID			ID of the gene that produced the transcript (parent gene)
nodeID			ID of the transcriptional event (either transcript name or the name of CCRE the transcript is assigned to)
clusterID		ID of the cluster the transcript is assigned to
geneName		name of the parent gene
linkageCluster		Linkage cluster of the parent gene, according to Desjardins et al. 2013
recombinationRate	Recombination rate of the parent gene, according to Desjardins et al. 2013
stageSexBias		Presence or absence of sex-bias at any of the five developmental stages. "f" indicates significant female bias, "m" significant male bias and "." no significant sex-bias at the given stage. Stages are ordered from early (left) to late (right). 
stageBias		Presence of up or downregulation at the given stage. "+" indicates upregulation, "-" downregulation, "." no significant changes. Stages are ordered from early (left) to late (right). 
strataMetazoa		Stratum assigned to the parent gene, as per Sackton, Werren, and Clark 2013
strataWasp		Stratum assigned to parent gene, including data from Lindsey et al. 2018
eventType		Whether the node represents a transcriptional event or a splicing event
Connectivity		Sum of all connection weights associated to the node, within its own cluster
hubScore		Connectivity weighted by clustering coefficient
