## Supplementary Files 1-5 for "Sex biased expression and co-expression networks in development, using the hymenopteran *Nasonia vitripennis*": S2_Readme.txt

clusterID			ID of the transcriptional cluster
Rows 2:11			See definitions in paper, section Multivariate Analysis of Network Parameters
stageSexBias			Presence or absence of sex-bias at any of the five developmental stages. "f" indicates significant female bias, "m" significant male bias and "." no significant sex-bias at the given stage. Stages are ordered from early (left) to late (right). 
stageBias			Presence of up or downregulation at the given stage. "+" indicates upregulation, "-" downregulation, "." no significant changes. Stages are ordered from early (left) to late (right). 
DifferentiallyCorrelated	Stage at which the cluster shows sex-bias in correlation, and direction.
