## Supplementary Files 1-5 for "Sex biased expression and co-expression networks in development, using the hymenopteran *Nasonia vitripennis*": S3_Readme.txt

transcriptID		ID of the transcript included in the CCRE
CCRE_ID			ID of the CCRE to which the transcripts are assigned
representativeNode	whether the expression values of this transcript used to represent the CCRE
